# Predicting peptide properties from mass spectrometry data using deep attention-based multitask network and uncertainty quantification

**DOI:** 10.1101/2024.08.21.609035

**Authors:** Usman Tariq, Fahad Saeed

## Abstract

Database search algorithms reduce the number of potential candidate peptides against which scoring needs to be performed using a single (i.e. mass) property for filtering. While useful, filtering based on one property may lead to exclusion of non-abundant spectra and uncharacterized peptides – potentially exacerbating the *streetlight* effect. Here we present *ProteoRift*, a novel attention and multitask deep-network, which can *predict* multiple peptide properties (length, missed cleavages, and modification status) directly from spectra. We demonstrate that *ProteoRift* can predict these properties with up to 97% accuracy resulting in search-space reduction by more than 90%. As a result, our end-to-end pipeline is shown to exhibit 8x to 12x speedups with peptide deduction accuracy comparable to algorithmic techniques. We also formulate two uncertainty estimation metrics, which can distinguish between in-distribution and out-of-distribution data (ROC-AUC 0.99) and predict high-scoring mass spectra against correct peptide (ROC-AUC 0.94). These models and metrics are integrated in an end-to-end ML pipeline available at https://github.com/pcdslab/ProteoRift.

## 1 Introduction

Reduction in search-space using peptide mass as a filter is one of the most fundamental techniques utilized to make database-search algorithms scalable. While mass is just one property of spectra and associated peptide, filtering does reduce unnecessary comparisons, minimizes the high-scoring miss-matches, and improves the search accuracy [1]. However, existing algorithms using precursor and fragment ion masses were not designed to handle the complexity of multiple non-model biological entities – such as meta-proteomics investigations [2, 3, 4]. This continuous nature of the mass filter which when used for meta-proteomics data analysis, leads to either large number of false positive peptides expelled from further analysis – or unreasonable search-times because of very large and redundant databases [4, 5, 6] – or at worst both. The mass spectrometry (MS) community is aware of unidentified peptides that do not get “matched” even when in the database, resulting in the development of “open-search” [7] and “hybrid-search” mechanisms [8, 9]. In “open-search” methods, the mass filter is increased significantly resulting in large number of candidate peptides included in the scoring which would otherwise may get missed. Methods such as MSFragger [10] have improved on the identification of the peptides, but execution of search-space restriction remain fundamentally the same i.e., mass of the spectra and corresponding peptides. However, among other factors, combination of technological limitations has resulted in inaccuracies including misidentification/no-identification of peptides [11], inconsistencies between search engines [12, 13, 2, 11, 14, 15, 16], tendency to identify abundant peptides [17, 13] - leading to the so-called street-light effect [17, 18, 19]. Unidentified peptides, and unaccounted post-translational modifications has led to multiple calls [17, 18, 20] by scientists to device methods to study understudied and non-abundant proteins associated with human aliments [21] including rare diseases [22], neurological disorders [23, 24, 25], and cancers [26, 27] - illustrates urgency and importance of the problem.

Advances in machine-learning (ML) models have made it possible to develop more accurate and deeper pipelines for MS data analysis [13, 28, 12, 29, 30, 31]. While it is widely believed that machine-learning models can outperform traditional (algorithmic) database and denovo searches, most new models have focused on scoring functions [32]. Routine usage of these models remains challenging [33] because ML scoring functions are inserted in the “algorithmic” workflow resulting in peptides similar to their algorithmic counterparts. Further, reluctance of systems biology labs to incorporate ML models influenced by multifaceted factors [34, 35, 36] such as black-box nature of most ML models, and the (less) confidence/certainty of the deduced peptides by the ML model are also contributing factor for this dearth. To reap the benefits of advancement in machine-learning technology, components, other than scoring, needed to perform MS data search must be developed to transpire an end-to-end ML pipeline.

In this paper, we designed and developed a novel deep-learning model to predict peptide features (peptide length, missed cleavages, and modifications) directly from spectra embedding – before any database-search. Our proposed network inputs: mass to charge ratio (m/z), intensity, and index values of spectra using the *attention network*, incorporate precursor mass and charge value of the spectrum to predict all the features simultaneously using the *multitasking* technique. Theoretical analysis shows that successfully applying filters reduces the search space size by more than 90% (Supplementary Figure 1) which is then demonstrated with experiments using real-world data (Fig. 3). Our extensive experimentation with open data demonstrates that the proposed model can predict peptide lengths by up to 92% precision, missed cleavages by 95% precision, and modifications status by up to 97% precision. We further retrained our ML model Specollate by incorporating newly designed filters in the ML workflow. These ML filtering models when integrated with database-search engine leads to 8x – 12x speedup while maintaining peptide accuracies at par with traditional highly successful search engines such as Crux or MSFragger. Lastly, we developed three novel MS-specific metrics to quantify the uncertainty associated with spectra embeddings, and their inferred peptides using re-trained SpeCollate [37] model. Our results demonstrate that the proposed metrics can distinguish in-distribution and out-of-distribution data (ROC-AUC 0.99) and predict high-scoring mass spectra against the correct peptide (ROC-AUC 0.94). These measures provide insight into the model’s stability, confidence, and data representation ability, aiming to empower confident usage of ML tools in systems biology settings.

## 2 Results

### 2.1 Method Overview

*ProteoRift* is a deep attention-based multitask network that enables prediction of peptide properties directly from the spectra. To show case the power of predictive analytics, we developed an end-to-end database search pipeline shown in Fig. 1 which enables comparison of results directly to traditional algorithms such as Tide and MSFragger. We designed and developed different stages of ML end-to-end pipeline for peptide deduction (Fig. 1). In the first step, attention-based network generates spectra, and peptide embeddings. Thereafter, a multi-task network is used to predict peptide-length, missed cleavages, and modifications status directly from spectra *before* any peptide deduction. The second step consists of binning spectra, and peptides using the predictions from the first step. These batches are then used for performing database-search for peptide deduction using spectra and peptide in single batch i.e., no intra-batch-deductions are performed. Provided that the first step resulted in a correct prediction, the peptides and the spectra that are *suppose* to match would be in the same batches. The third step is to develop novel metrics to quantify the uncertainty associated with spectra embeddings and peptide deduction. These confidence metrics are reported back with the results of the spectra-to-peptide matches. This complete end-to-end ML pipeline is then used for all experimentation as well as comparison with existing methods. To the best of our knowledge, we demonstrate first *end-to-end* ML pipeline that predicts peptide properties directly from the spectra, and enable more scalable, accurate, and trustworthy peptide deductions.

**Figure 1:**
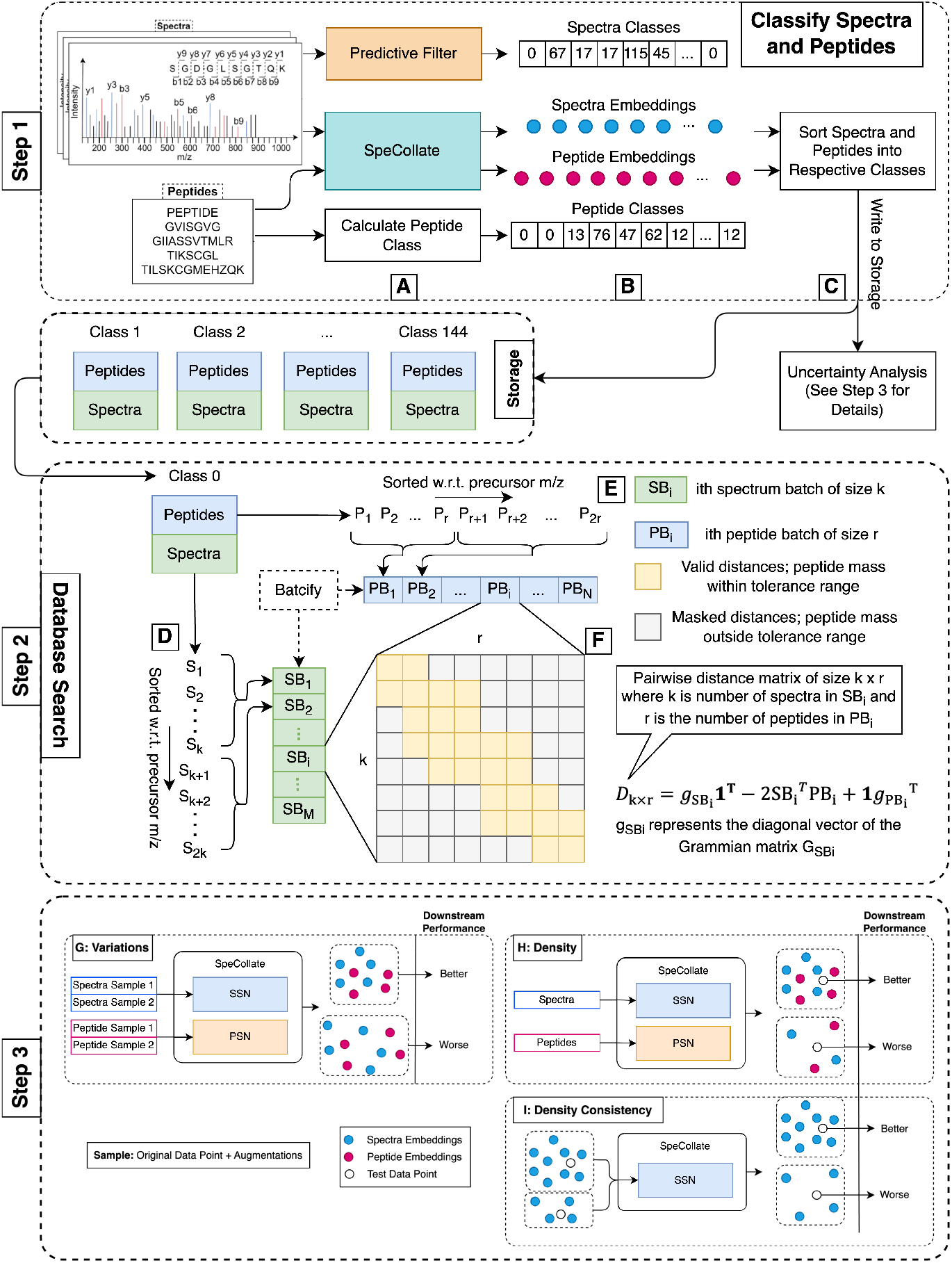
Filtered Meta Proteomics database search pipeline. **Step 1** (A) Spectra and peptide embeddings are generated using SpeCollate while the predictive model assigns a class (out of 144) for each spectrum. (B) Class for a peptide can be calculated from the amino acid sequence. (C) Store spectra-peptide classes. **Step 2**: (D, E) Batchify spectra and peptides to fit them in server memory. (F) Calculate distance matrix of spectra and peptide embeddings in a given batch. **Step 3**: (G) Per-feature sample variations and their correlation with downstream performance shown. A sample, the original data point plus augmentation, has lower feature variations when the middle is generally more confident about the original data point. (H) The density of training data around a given test data point indicates higher model confidence and hence improved downstream performance. (I) Density consistency measures how consistent the input data density is with the embedding density. This feature is important in estimating how well the model is able to capture the data structure and relations.

### 2.2 Experimental Setup Overview

We constructed Pride Archive (PXD) datasets for our experimentation and evaluation for proteomics search: PXD000612 [38], PXD001468 [39], and PXD009861 [40]. When extending our experiments to Meta-Proteomics database searches, we use the RefUP++ database [41], encompassing 2,259 genera. To gain insights into our predictive models we performed experiments for 3 distinct sets. The first set of experiments were performed for evaluating the results of the predictive model i.e. how accurate is our developed *ProteoRift* model is when trying to predict the length, missed cleavages and modifications. The second set of experiments were performed for the database search that is batched according to the model predictions. This set of experiments informs us about the performance of the end-to-end pipeline as compared to existing algorithmic techniques. Furthermore, these experiments inform us about the potential search-space and processing time reduction. The last set of experiments are completed to assess the confidence of the ML database search embedding, and apprises us about the confidence metrics when compared with the ground truth data sets.

#### 2.2.1 Runtime environment

The experiments are performed using the PyTorch framework 1.13.0 and Python 3.10.8. For fast training, we utilized NVIDIA V100s (32 GB SMX2) GPUs with 32GB of memory provided by Expanse SDSC supercomputer installed in nodes with up to 4 GPU nodes, 40 CPU cores, and 384 GBs of memory (10 CPU cores and 96 GBs per GPU node). In addition, NVIDIA A6000 GPUs with 48GBs of memory were utilized as part of our in-house cluster (dragon).

#### 2.2.2 Train, Test, Validation Datasets

The model is trained using high-quality spectra with selected label peptides. Three datasets i.e., NIST [42], MassIVE [43], and DeepNovo [44] datasets are used. The spectral libraries are preprocessed to extract the MS/MS spectra along with the target labels from the corresponding peptide i.e., peptide sequence length, missed cleavages, and modification status. Peptide sequence length is calculated as the number of amino acids in the peptide regardless of the modifications on any amino acid. Missed cleavages are calculated as the number of K and R amino acids within the sequence except if followed by P or when at the end of the peptide sequence. Modification status is calculated as true or false depending on whether there is a dynamic modification (one or more) on any number of amino acids. Carbamidomethylation (CAM) on amino acid C is not considered a modification for our data. We obtained ∼2.7M spectra with known peptide labels out of which ∼.37M were modified, and the rest were unmodified. The dataset contained ∼1.7M spectra with 2+ precursor charge and ∼0.5M spectra with 3+ precursor charge. In terms of missed cleavages, the training dataset contains ∼2M, ∼0.7M, and ∼50k spectra with zero, one, and two missed cleavages respectively. Train/Validation/Test split of 0.7/0.2/0.1 was used for training the network and tuning hyperparameters. Now, our dataset only contains oxidation modifications. The precursor charge value of the spectra was restricted to a maximum of 3+. The distribution of length missed cleavages, and modification status is shown in supplementary figure 3.

### 2.3 Accuracy of the Predictive power of *ProteoRift*

On the test dataset, we measure the precision values of all classes for each head.

#### 2.3.1 Performance of length head

Our model predicts the length with high precision for smaller lengths which then gradually drops for large length values Fig. 2 (top left). This is to be expected because of limited data available for us train the models for larger length peptides. The fact that the total precision value is *>* 90% is observed for most length within ± 1 demonstrates that the head is sufficiently trained and predicts lengths with high precision. In light of these results, we consider *predicted*_*length* = *actual*_*length* ± 1 to be accurate for any subsequent experiments and the model.

**Figure 2:**
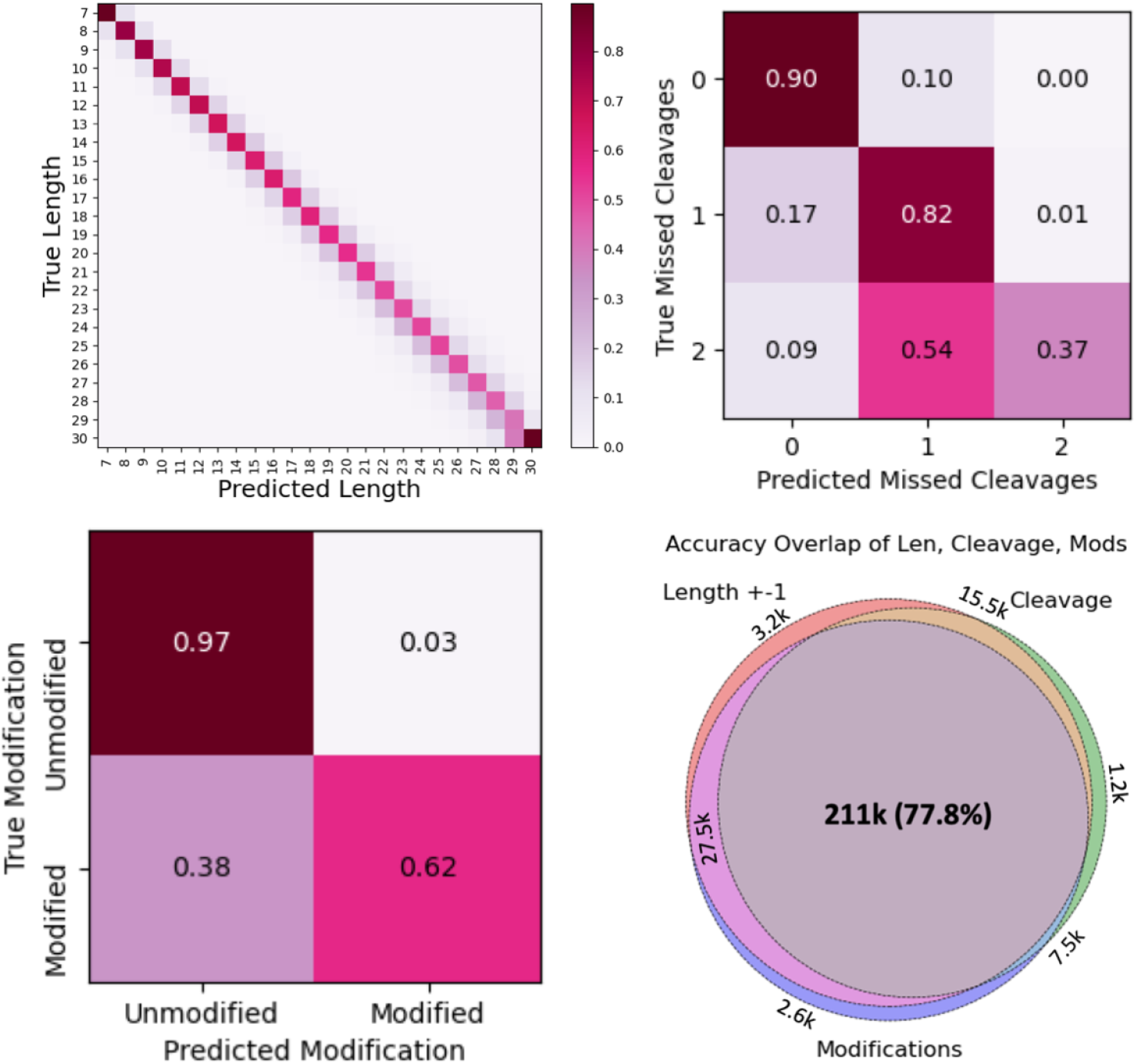
Confusion matrix of peptide length, missed cleavages, and modification status predictions. For length prediction, precision is high for smaller lengths which gradually drops for larger length values, since only fewer number of training samples are available at larger lengths (similar to in real-life MS experiments). When peptides within *±* 1 of the predicted length are considered, precision value is > 90% for most lengths. The illustrated Venn diagram depicts the correct predictions of all three heads. ProteoRift accurately predicts all three properties 77.8% of the time. Note that for length prediction, we use the recommended error tolerance of *±*1 for optimal performance.

**Figure 3:**
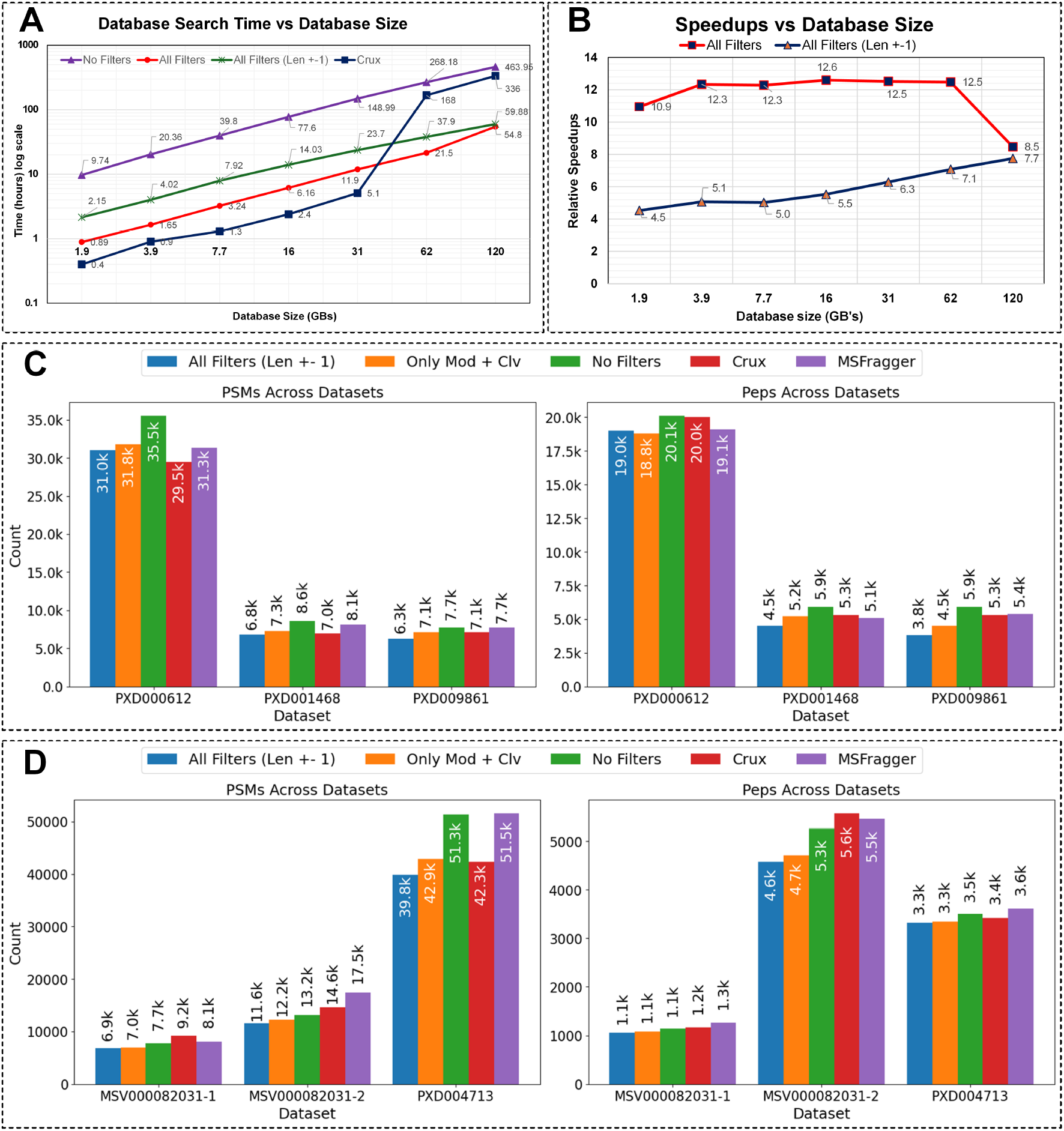
**A and B**: Time of execution and speedup analysis with different filter configurations and increasing database size for a Meta-Proteomics database. The search is performed against a meta-proteomics database published by Proteostorm (RefUP++ database [41]). The mass spectra from the msv000082031 spectral dataset was used. When all filters are applied we obtain up to 12x speedup while 8x speedup when enabling the length error margin of *±* 1. **C and D**: Comparison of the number of PSMs and peptides identified under 1% FDR against SpeCollate with and without length filter. Results for Crux and MSFragger are also illustrated. The number of peptides identified when applying predictive filter is similar to when no filters are applied, while obtaining significant speedup.

#### 2.3.2 Performance of cleavage head

Our experiments also demonstrate that head to predict cleavage also works with high precision (Fig. 2 (top right)). As shown in the figure, the 0 and 1 missed cleavages are predicted with 90% and 82% respectively. However, the precision drops rapidly for 2 missed cleavages. This is again to be expected since 2 missed cleavages are rare in real-world data which is also apparent in our training data sets. For our experiments, we ensured that peptide database is generated with an equal portion of 2 missed cleavage peptides, and therefore the effective precision in real-world setting would be much higher providing considerable benefit in terms of reducing search space.

#### 2.3.3 Performance of modification-status head

Our model predicts with exceptional performance of more than 97% precision for unmodified spectra as shown in Fig 2 (bottom left). While the performance for much difficult problem of modification status is close to 62%, it is much better than random chance and will result in reduction in the search-space. This behavior is also expected since the number of modifications are very large in number, leading to explosion in possible modification patterns in the spectra. While the model did learn, and performed better than random for modification status on unseen data; more data may lead to better performance in the future.

#### 2.3.4 Overall predictive performance

The average performance of three heads will determine the accuracy of the overall predictive model and it effect on the search-space. We perform the aggregate performance evaluation of the model by measuring the total number of correct measurements for all heads. As shown in Fig. 2 (bottom right), we are able to accurately predict all three properties 77.8% of the time.

### 2.4 Effect of predicted search-space on peptide deduction completed using end-to-end ML database-search pipeline

These set of experiments were completed to estimate the effect of the filters on the search-space as well as the accuracy of the peptides deduced using modified search-space. For our predictive filtering to be effective, the peptide deductions on modified search-spaces must be at par with other algorithmic techniques. To perform comprehensive evaluation of the effect of the predictive filtering on peptide deduction quality and execution time, we incorporate various permutations of the filter. This predictive filter configuration includes: 1) applying all filters simultaneously, 2) selectively applying missed cleavages and modification filters, and (3) implementing all filters in conjunction with a relaxed length filter with a margin of ±1.

#### 2.4.1 Reduction in search-space and speed up analysis

The first thing we needed to establish was if predictive filtering effectively reduced the search-space when deducing peptide from database-search. The search is performed against a meta-proteomics database published by Proteostorm. The mass spectra used were from the msv000082031 spectral dataset. For the Meta-Proteomics search experiments, the database size varies from 2 GB to 120 GB. As shown in Fig. 3A and B, the speedup after using filters increases gradually as the database size increases. This is likely due to the reduced number of out-of-core computations when additional filters are applied leading to a more optimal memory access pattern. While we see that crux, which is highly optimized piece of code, better than our unoptimized search-code, the advantage disappears as the size of the database increases. For any optimized code we must expect better speedups than are reported here. In general, our experiments demonstrate that when all filters are applied, we obtain up to 8x to 12x speedups. Speedups of up to 7.7x are obtained when when enabling the length error margin of ±1. This is predominantly attributed to a significant reduction in out-of-core computations, a frequent computational bottleneck in large-scale database searches. Since predictive filtering does result in smaller search-space and better execution times especially for larger databases, we next compare the peptide deduction results with existing state-of-the-art algorithmic techniques.

#### 2.4.2 Proteomics data sets

Since predictive filtering model can be used with any database search-engine we also wanted to assess its effectiveness when deployed for proteomics experiments. For varying filter configurations we evaluated three distinct datasets, namely PXD000612 [38], PXD001468 [39], and PXD009861 [40]. In these trials, we consider two primary metrics: the number of identified peptides under a 1% False Discovery Rate (FDR) and the resulting speedup of the database search process. As shown in Fig. 3C, the PSM results with no-filter (Specollate) is better than both Crux and MSFragger for PXD000612 and PXD001468 and comparable to MSFragger for PXD009861. For PXD000612, Mod+cleavage filter performs better than both Crux and MSFragger for PSM and performs comparably for the other two data sets. When all filters are used, which is arguably the most stringent settings, the PSM identified are still comparable while giving 8x to 12x speedups as compared to non-filtered settings. The number of peptides identified when using mod+cleavage filter is comparable to MSFragger and Crux for all data sets.

#### 2.4.3 Meta-proteomics data sets

When extending our experiments to Meta-Proteomics database searches, we use the RefUP++ database [41], encompassing 2,259 genera. We analyze the number of peptides identified (at 1% FDR) with different configurations for the Meta-Proteomics database. Our results shown in Fig. 3D, demonstrates that number of peptide-to-spectrum matches, and peptides identified are comparable to MSFragger and Crux for msv000082031-1, msv000082031-2 and PXD004713. For PXD004713 the number of PSM’s using filtering technique surpasses Crux when only modification and cleavage predictive filtering is used. While the number of PSM is higher for MSFragger, the number of unique peptides identified are comparable when predictive filtering is used. Lastly, the results are extremely encouraging considering that the number of peptides identified with filtered subsets is like when no filter is applied highlighting that our predictive filters can provide significant speedup while having a negligible effect on result quality – while exhibiting impressive 12x speed ups. Note that this improvement is a vast advancement over the speedup obtained by solely applying the precursor mass filter, further illustrating the value of predictive filtering.

### 2.5 Predictive power of confidence and uncertainty metrics and their correlation with real-world datasets

The utility of database-search ML models might be subject to various degrees of vagueness influenced by multifaceted factors [34, 35, 36]. This imprecision encapsulates two key dimensions: *aleatoric* and *epistemic* [45]. Aleatoric imprecision originates from the inherent noise and stochasticity in the data, whereas epistemic imprecision mirrors the model’s limitations, signifying the unlearned or unknown aspects within the model’s purview. Such uncertainty estimation enables the reliability of the model’s predictions. For example, elevated uncertainty in peptide or spectrum embeddings may signal model ambiguity in their representations, potentially indicating a misaligned peptide-spectrum match. Further, such uncertainty quantification provides insights into embedding areas that can be improved either by further training or acquiring more quality or diverse data. For example, higher epistemic uncertainty can indicate regions in the input space where the model is *under-confident* due to a lack of sufficient data. This can point out opportunities for further data collection or model improvement. Similarly, higher aleatoric uncertainty can point out the need for higher-quality data or preprocessing for noise removal. Most importantly, uncertainty estimation allows for risk-aware decision-making. In high-stakes applications such as clinical proteomics — where incorrect peptide identification could lead to misleading conclusions — being aware of the uncertainty associated with each prediction can help avoid potentially costly or harmful decisions.

Our objective is to design uncertainty metrics that can be used to assess the confidence in the peptide deduction i.e., how confident the scientist should be in the inference especially when the identified peptide is novel. We estimate the aleatoric and epistemic uncertainty of SpeCollate’s embeddings using our proposed metrics and analyze how they can inform us about the model’s performance and its output confidence. To this end, we developed three different metrics for estimating the uncertainty of the embeddings: 1) We assess the certainty of embedding location by introducing controlled augmentations to the input spectra and then measuring the variation in the output embeddings. 2) The density of the training data around each embedding is measured using a von Mises-Fisher (vMF) Mixture Model to determine how much data the model has seen around a given data point. 3) We evaluate whether the density is consistent between the input spectra and their embeddings, indicating whether the model can maintain data structure and relationships.

#### 2.5.1 Aleatoric Uncertainty

To determine how well our metrics estimate aleatoric uncertainty for a given embedding, we use a mass spectrometry dataset that is similar to SpeCollate’s training data i.e., ProteomeTools HCD data [46]. We want to see how well these metrics can predict the downstream performance of the in-distribution embeddings e.g., in our case, how well the mass spectra embeddings can be identified in a peptide database search. Ideally, higher confidence along with a higher ranking of the spectrum would indicate the model is certain about its decision. To quantify this, we rank peptide embeddings from the human peptide database against each spectrum. A true label is assigned to a spectrum if the correct peptide is ranked the highest. The label definition is given below:

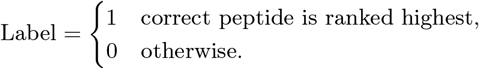

We train a gradient-boosting decision tree (GBDT) model with per-sample feature variations (variations for short), density, and density consistency as input and the above-described labels as output. We perform hyperparameter tuning for the max depth, learning rate, and number of estimators. The optimal values selected are shown in supplementary table 1.

Detailed analysis of the metrics performance is given in table 1. As can be seen in the table, the per-sample feature variation (variation) metric yielded a promising ROC-AUC score of 0.84, surpassing the other individual metrics. This finding illustrates that the model’s confidence in an example’s embedding position is a crucial indicator for identifying high-quality spectra. Combining the variations and density metrics resulted in a significant enhancement in all performance metrics emphasizing that the two metrics combined capture different aspects of uncertainty providing overall better performance in retrieving high-quality examples. The combination of all three metrics yielded the best performance across all metrics, with a ROC-AUC score of 0.94. This reveals that a holistic view, encompassing variations in embedding, data distribution around the embedding, and consistency between input and embedding densities, provides the most potent approach for identifying high-quality spectra.

**Table 1:**
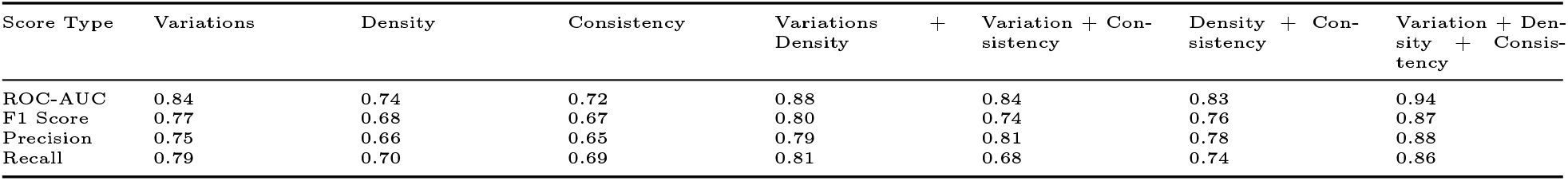
Performance of our uncertainty metrics on retrieving high-quality examples from in-distribution data. The values indicate that combining multiple metrics improves the ability to detect high-quality examples significantly with the ROC-AUC value of 0.94.

#### 2.5.2 Epistemic Uncertainty

To estimate epistemic uncertainty, we aim to classify in-distribution and out-of-distribution data points using our uncertainty metrics. For in-distribution and out-of-distribution data, we use ProteomeTools HCD and CID datasets. As SpeCollate is solely trained on HCD data, CID data is an obvious choice for out-of-distribution examples to validate our uncertainty metrics. We assign the following labels to our curated dataset with nearly 1 million examples for each class:

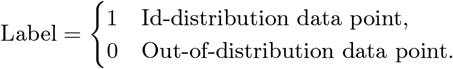

We train a GBDT model on this data with a 70-20-10 split of train validation and test subsets. The optimal parameters selected after hyperparameter tuning are given in supplementary table 2.

The standout performance of the consistency metric (ROC-AUC of 0.99) reinforces its innovative design. Assessing the alignment between the density of input data, and embeddings offers insight into the consistency of the model’s behavior, providing a crucial aspect of uncertainty evaluation. Detailed results are given in table 2.

**Table 2:**
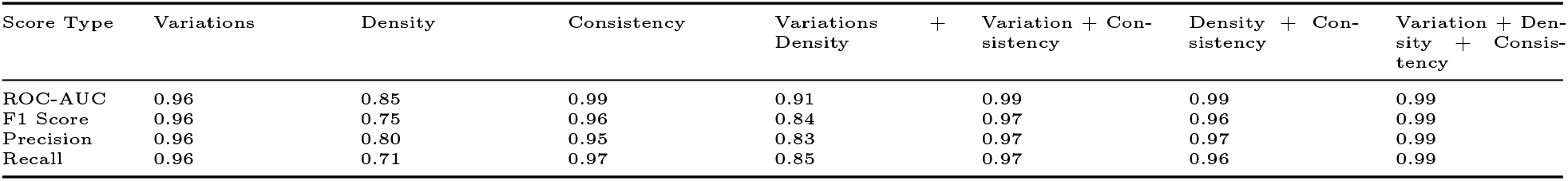
Performance of our uncertainty metrics on retrieving in-distribution data from a mix of in-distribution and out-of-distribution examples. The values indicate that density consistency is the major contributor towards distinguishing between the two classes with an ROC-AUC of 0.99.

To further validate our results, we perform a database search on both in-distribution and out-of-distribution datasets. We selected three datasets where each dataset is a mixture of in-distribution and out-of-distribution data. Confusion matrices are calculated to determine what percentage of in-distribution predicted spectra match with the correct peptide. As shown in Figure 4, the predicted in-distribution data points perform vastly better than the predicted out-of-distribution data with an accuracy of up to 96.4%.

**Figure 4:**
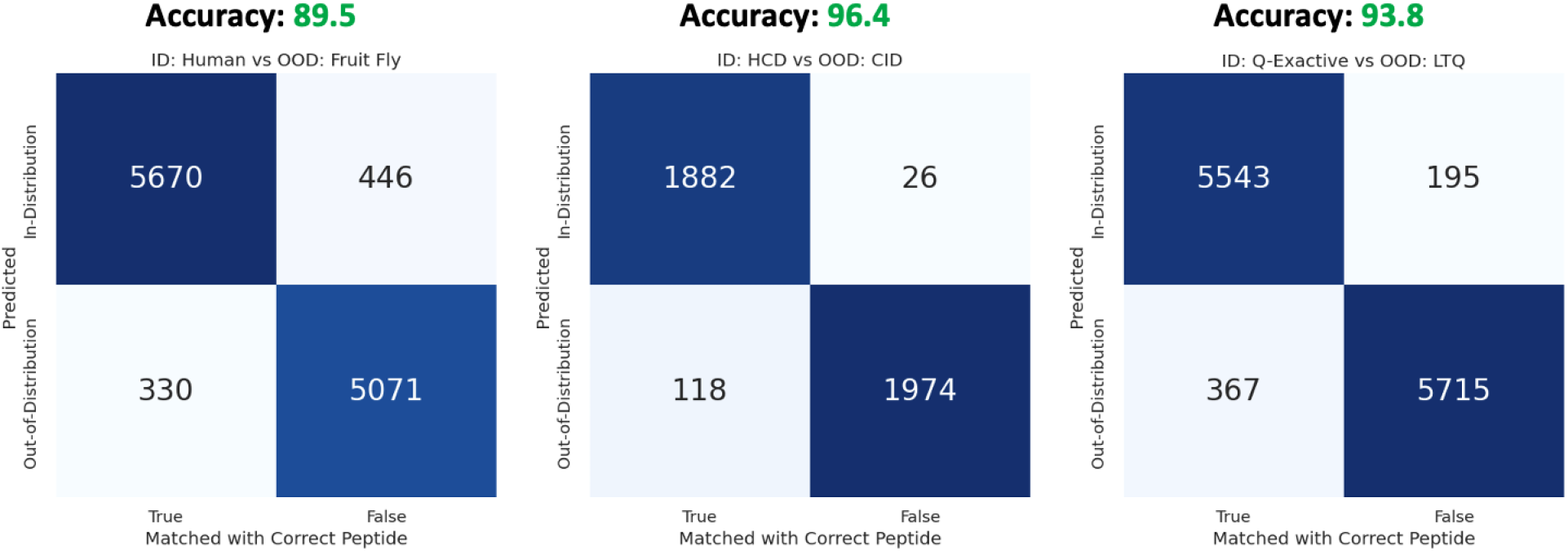
Peptide identification accuracy of spectra predicted in-distribution vs out-of-distribution. Three different combinations of in-distribution and out-of-distribution datasets are used to compare the downstream performance with the predicted accuracy. For Human vs Fruitfly, we use datasets PXD009861 and PXD030281. For HCD vs CID, we use ProteomeTools HCD and CID libraries. For Q-Exactive vs LTQ, we use datasets PXD000612 and PXD000125. As can be seen in the figure above, the in-distribution data points can be predicted with all degree of accuracy with all different data sets

## 3 Discussion

*ProteoRift* is a machine-learning method that can predict multiple peptide properties (length, missed cleavages, and modification status) directly from spectra. We demonstrated that these properties when used for search-space reduction, results in superior speeds (8x to 12x faster) and comparable results for both proteomics, and meta-proteomics experiments. Given that different properties of peptides and the spectra can be used for search-space reduction, the usage of mass-only filtering by existing methodologies is a bottleneck, especially for searches that involve proteogenomics, meta-proteomics or non-model organisms. In this paper we also introduced certainty metrics which will enable confidence estimation of the results especially for identification of novel and uncommon peptides – potentially reducing the skepticism of results obtained using black-box machine-learning models.

ProteoRift is currently a prune-and-search method which can result in reduction in accuracy with successive pruning attempts. Accumulation of more filtering decreases the search-space, and results in decreased false-positives. However, this also results in decrease in the number of identified peptides due to summation of the errors by three different heads. This model prediction error can be attributed to the imbalance in data (supplementary figure 3) where the distribution of features is highly skewed and some labels have very few training examples. ProteoRift is a supervised learning technique and the amount of labelled data available is one of the limitations of the method. Therefore, the accuracy of the peptide properties achieved using ProteoRift, and subsequent peptides deduced, are limited by labelled data that is currently in short supply.

We believe that machine-learning tools are the enablers of new exciting science. However, the research community needs more synergistic efforts in integrating ML scoring function with existing or all-ML pipelines. Additionally, improved scoring techniques using machine-learning, for both denovo and database search, are steps in the right directions, other parts of the workflow are equally important for discovery of non-abundant peptides and proteins and is part of our future work. To this end, we have demonstrated an end-to-end pipeline for peptide deduction, and integration of uncertainty measures will enable scientific investigations and adaptation of ML tools in this domain. Such ML computational infrastructure will potentially reduce the street light effect, resulting in discovery of uncommon peptides and proteins – related to human disease and health.

## 4 Methods

In this section, we will present the design and implementation of our end-to-end peptide database search pipeline with predictive filtering and uncertainty analysis. We design a deep attention-based multitask network (ProteoRift). Four significant contributions of this work is the design and development of: 1) attention-based network to generate spectra and peptide embeddings, 2) multi-task network to predict peptide-length, missed cleavages, and modifications status directly from spectra, 3) novel metrics to quantify the uncertainty associated with spectra embeddings, and 4) predictive filtering based peptide database search that takes into account the confidence of spectra embeddings.

### 4.1 *ProteoRift* : A machine-learning model to predict peptide properties directly from the MS spectra

#### 4.1.1 Spectra and Peptide Embeddings

Our first challenge is to translate spectra and peptides’ high-dimensional sparse vectors into embeddings that can place semantically similar spectra and peptide inputs close together in the embedding space.

We have designed and developed a embedding network shown in Fig 5. The network uses self-attention layers to embed mass spectra. Using self-attention layers for embedding mass spectra is useful in capturing long-distance relations [47] between peaks e.g., complimentary b and y peaks as well as short-distance relations e.g., a neutral loss or isotopic peak (supplementary figure 2). For successfully capturing all the information in the spectrum, m/z, and intensity sequences are first discretized i.e., m/z values are binned in 0.1 Da sized bins, and intensity values are discretized between 0-1000 before passing them through the embedding layers. Since attention layers processes the entire sequence at the same time, the sequence information needs to be encoded separately. For this purpose, we use the sinusoidal harmonics as described in [47]. The m/z and intensity embedding as well as the sequence encoding are of the same size 256 and are added together before being processed by the attention layer.

**Figure 5:**
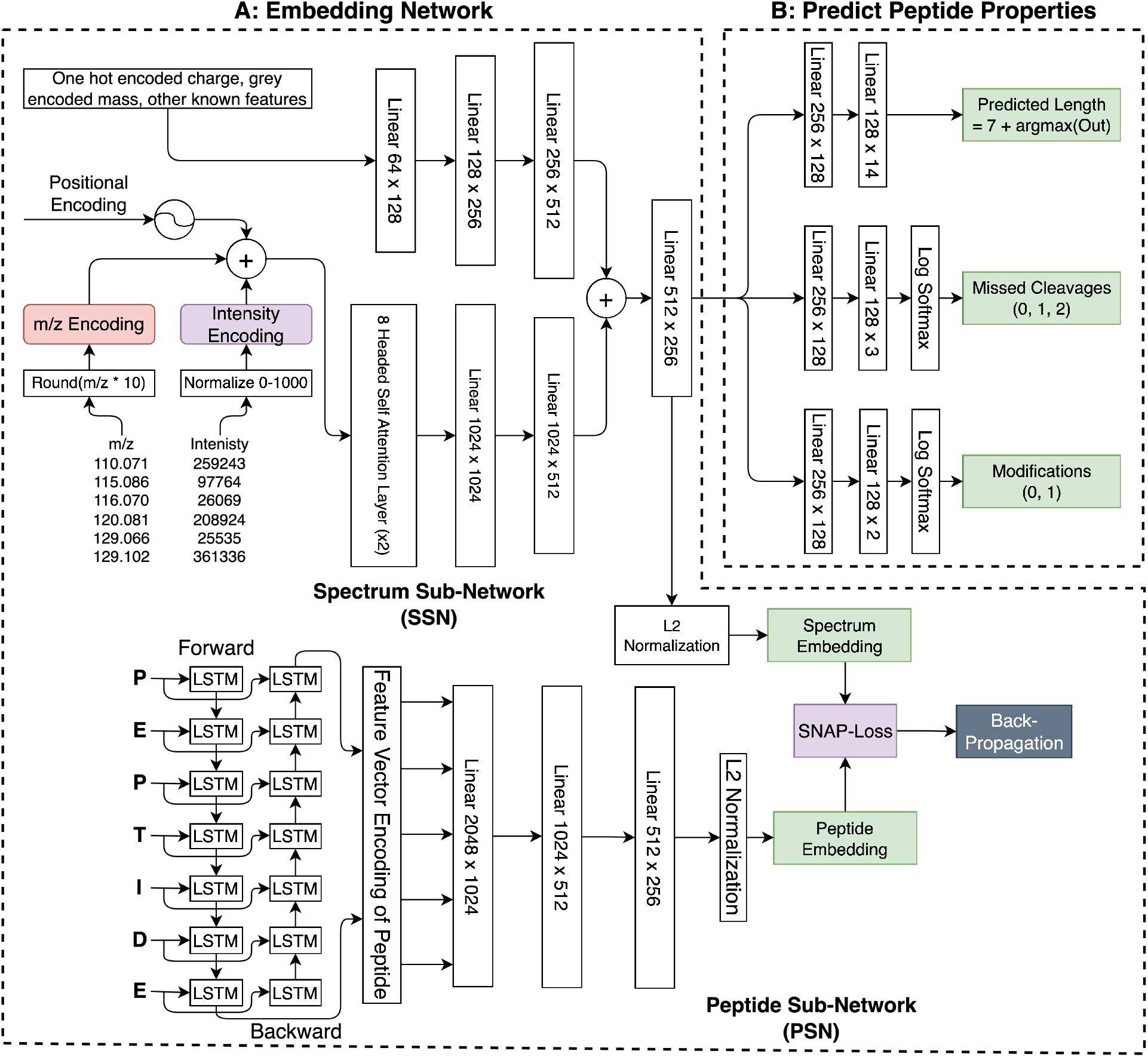
Network architecture for generating spectra and peptide embeddings. In SSN, spectra m/z and intensity values are treated as two separate sequences and embedded using separate embedding layers. Sinusoidal positional encoding is also added to the embeddings to maintain sequence order information. An intermediate spectrum feature vector is added with charge and mass feature vectors and passed through a fully connected layer of 512 x 256. In PSN, peptide embeddings are generated using two Bi-LSMT layers followed by three fully connected layers. From the spectrum embeddings, three branches of fully connected layers predict the peptide length, missed cleavages, and modifications as shown in part B. All the output branches calculate loss separately using the LogSoftmax function and a weighted sum of three losses is calculated before calculating gradients.

Next, we use two sixteen-headed attention layers to capture the sequence dependencies in the spectrum. The attention layers are followed by two fully connected layers of size 2048 x 1024 and 1024 x 512. The feature vector of mass and charge values is added to the intermediate output after the second fully connected layer. The combined feature vector is passed through another fully connected layer to generate spectrum embedding of length 256.

Peptide embedding are generated using 2 Bi-Directional Long Short-Term Memory (Bi-LSTM) networks. Amino acid level embedding are generated using an embedding layer, and fed to the Bi-LSTM which generates output vectors of size 2048 per peptide. Note that only the final amino acid output is kept and fed to the following layers. Next, we use two fully connected layers of size 1024, 512, and 256. The final layer is preceded by L2 normalization to generate normalized vectors of length 256 for each peptide.

#### 4.1.2 Predicting peptide properties

From the spectrum embeddings, three branches of fully connected layers predict the peptide length, missed cleavages, and modifications as shown in Figure 5 B. For the length prediction, the two fully connected layers of 256 x 128 and 128 x 24 are used. The output layers are of length 24 because we predict lengths from 7 - 30 AAs. The missed cleavage branch contains two fully connected layers of size 256 x 128 and 128 x 3 as we only consider max of two missed cleavages. Finally, the modification branch consists of two fully connected layers of size 256 x 128 and 128 x 2. For the current version of ProteoRift, we only predict whether the spectrum is modified or not. In future versions, we will aim to predict the exact number of modifications as well as the specific known modifications that exist in the spectrum. All the output branches calculate loss separately using the LogSoftmax function and a weighted sum of three losses is calculated before calculating gradients.

#### 4.1.3 Network Training and Evaluation

The training process begins with a forward pass of the batch containing encoded spectra. Three sets of labels are used i.e., peptide length, missed cleavages, and modifications status. The network generates three separate outputs for each feature respectively which are treated as multi-class classification. The cross-entropy loss function is applied to each feature separately to calculate three loss values. Before the backpropagation is performed, a weighted sum of the three losses is calculated where the weights are determined through a randomized search. Adam optimizer is used for gradient updates with a learning rate of 0.0001 and weight decay of 0.00005. A dropout of 0.3 is used after the attention, and the fully connected layers during the training step. The dropout layers are disabled during the evaluation step. The network is trained for 500 epochs and achieves 92% peptide length precision, 97% missed cleavage precision, and 97% modification precision as shown in supplementary Fig. 4.

### 4.2 Implementing End-to-End Database Search with Predictive Filters

To implement the database search with the new filters we first pre-assign the database peptides and the experimental spectra into one of the 144 (7-30 peptide length ×0, 1, or 2 missed cleavages ×2 modifications statuses) classes based on their features (predicted from the ML filter). For the database peptides, the features are known and hence each peptide gets classified into a non-overlapping class. Spectra are classified based on their predicted features using the ProteoRift model. A spectrum from a given class only gets searched against peptides in the same class. Since each class is non-overlapping, 144 parallel searches can be performed simultaneously. The general filtered database search flow is shown in Figure 1 step 2.

In the first step, spectra and peptides are indexed into their respective classes based on their properties. Each spectrum and peptide get assigned to a unique class. Spectra and peptides within each class are batched. As shown in Figure 1 step 2, spectra in a given class e.g., 7-1-0 (Peptide length: 7, 1 missed cleavage, unmodified) are sorted with respect to the precursor mass, and split into batches (SB) of size 1024. Similarly, peptides are sorted with respect to their precursor mass, and candidate peptide lists are generated for each SB. To generate a peptide list for a given SB, peptides in the mass range [MinSB - PreTol: MaxSB + PreTol] are obtained. Here, MinSB and MaxSB are the minimum and maximum precursor masses in a given SB respectively. PreTol is the user-specified precursor tolerance configuration. Next, each peptide list is split into batches (PB) of 1024 peptides. The size can be adjusted according to the available memory. For each PB in a peptide list, a distance matrix against the SB is calculated. A mask is calculated for each spectrum where peptides outside the mass filter range are ignored. Next, the values in the distance matrix are sorted for each spectrum in increasing order of L2 distance, and k smallest distance peptides are kept for each spectrum as these are the highest-scoring peptides. The inverse of the L2 distance is reported as the match score. The process is repeated for each SB and results are written to a percolator [48] input (pin) file. A similar process is repeated for the decoy database for FDR analysis.

### 4.3 Uncertainty Analysis

In this section, we will discuss the proposed uncertainty measures and present their mathematical formulation.

#### 4.3.1 Per-Sample Feature Variation

The incorporation of augmentation techniques to estimate the sample variance is a method employed to quantify the level of certainty a model has regarding the position of its predicted embedding. The principle lies in the deliberate modification of each data point in multiple ways, e.g. random omission of a certain percentage of peaks or adjusting the charges associated with spectra. Sample feature variation is shown in Fig. 1 (step 3 part G).

Given an original spectrum *s*, we induce *n* augmentations thereby creating a set of altered spectra denoted by {*s*_1_, *s*_2_, …, *s*_*n*_}. Each derived spectrum embedding is a 256-dimensional vector. The variance, a statistical measure of the extent to which the data points deviate from their expected values is computed for each of these dimensions.

Mathematically, the sample variance *σ* can be defined as follows:

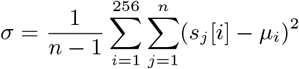

where *s*_*j*_ [*i*] denotes the *i*-th element in the *j*-th augmentation, *μ*_*i*_ is the mean of the elements for the *i*-th dimension, and *n* is the number of augmentations.

To compile these individual variances into a single value, a cumulative sum is calculated across all dimensions. This total sum, or sample variance, is indicative of the model’s confidence in the location of the predicted embedding; a smaller variance suggests a higher degree of certainty.

In essence, this technique provides valuable insights into the model’s confidence in its predictions, which can be crucial in the interpretation of the downstream performance. Only a trained model is required to estimate the variance.

#### 4.3.2 Density

The density of the embedding space around a spectrum embedding is the measure of how many training examples the model has seen close for a given example. We use the von Mises-Fisher distribution to estimate the density of the embedding space around a given data point. As the spectra and peptide embeddings generated by SpeCollate are L2-normalized, the von Mises-Fisher distribution, particularly when combined into a mixture model (vMFMM), is an ideal choice for modeling data due to its directional properties. Density estimation is visualized in Fig. 1 (Step 3 part H).

Assuming we have a set of N training examples {*s*_1_, *s*_2_, …, *s*_*N*_} and these are normalized to unit length; let’s say we fit a vMFMM to this data with K components. Each component k has a center *μ*_*k*_ and a concentration parameter *κ*_*k*_ and a weight *w*_*k*_. The density *p*(*s*) for any point *s* in this mixture model can be computed as:

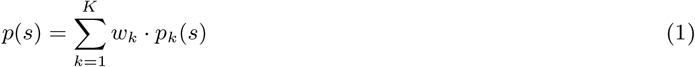

Where *p*_*k*_(*s*) is the probability density of spectrum embedding *s* in the *k*-th von Mises-Fisher component, defined as:

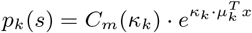

Where *C*_*m*_(*κ*) is the normalization constant in *m* dimensions, given by:

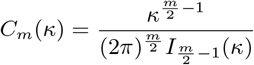

And 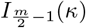 is the modified Bessel function of the first kind.

Therefore, to measure the density of training examples around a given point, we would compute *p*(*s*) using the above equations. This gives an idea of how familiar the model is with the region around a given embedding.

#### 4.3.3 Density Consistency

We propose a novel metric, herein referred to as the Density Consistency (DC), which is designed to assess the preservation of density consistency between the input data and the model’s output embeddings. The DC is a measure of a model’s ability to maintain the structural and relational characteristics of the data in its transformation to the embedding space. Density consistency is shown in Fig. 1 (Step 3 part I).

Given a data point *x* and its embedding *s* the DC is mathematically formulated as follows:

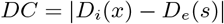

In this formula, *D*_*i*_(*x*) is the density of the input data around a given point *x*, which is estimated using domain-specific knowledge, and *D*_*e*_(*s*) is the density of the embeddings around a given point *s*, which is estimated using the von Mises-Fisher Mixture Model (vMFMM) as outlined in the previous section. The absolute value of the difference between these two densities yields the DC. In the case of our specific application, MS/MS spectra, the density of the input data *D*_*i*_(*x*) is calculated using the cosine similarity metric. This serves to gauge the similarity between pairs of input spectra. This metric, therefore, allows us to quantitatively evaluate the model’s performance in terms of its ability to maintain the integrity of data relationships and structure in the transformation from the input space to the embedding space. We argue that the lower the value of DC, the more effectively the model preserves the original data structure in its output embeddings.

## Code Availability

All code, associated models and weights are made available at: https://github.com/pcdslab/ProteoRift

## Data Availability

The datasets and database used in this study are publicly available from the mentioned respective data repositories. The experiment configuration files and raw results pertinent to the findings of this study are available from the corresponding author upon request.

## Acknowledgments

This research was supported by the NIGMS of the National Institutes of Health (NIH) under award number: R35GM153434. The authors were further supported by the National Science Foundations (NSF) under the award number: NSF OAC-2312599. The content is solely the responsibility of the authors and does not necessarily represent the official views of the National Institutes of Health and/or National Science Foundation.

The authors would like to thank Bilal Shabbir for editing the GitHub repository for wider dissemination. This work used the NSF Extreme Science and Engineering Discovery Environment (XSEDE) Supercomputers through allocations: TG-CCR150017 and TG-ASC200004.

## Supplementary Figures

**Supplementary Figure 1:**
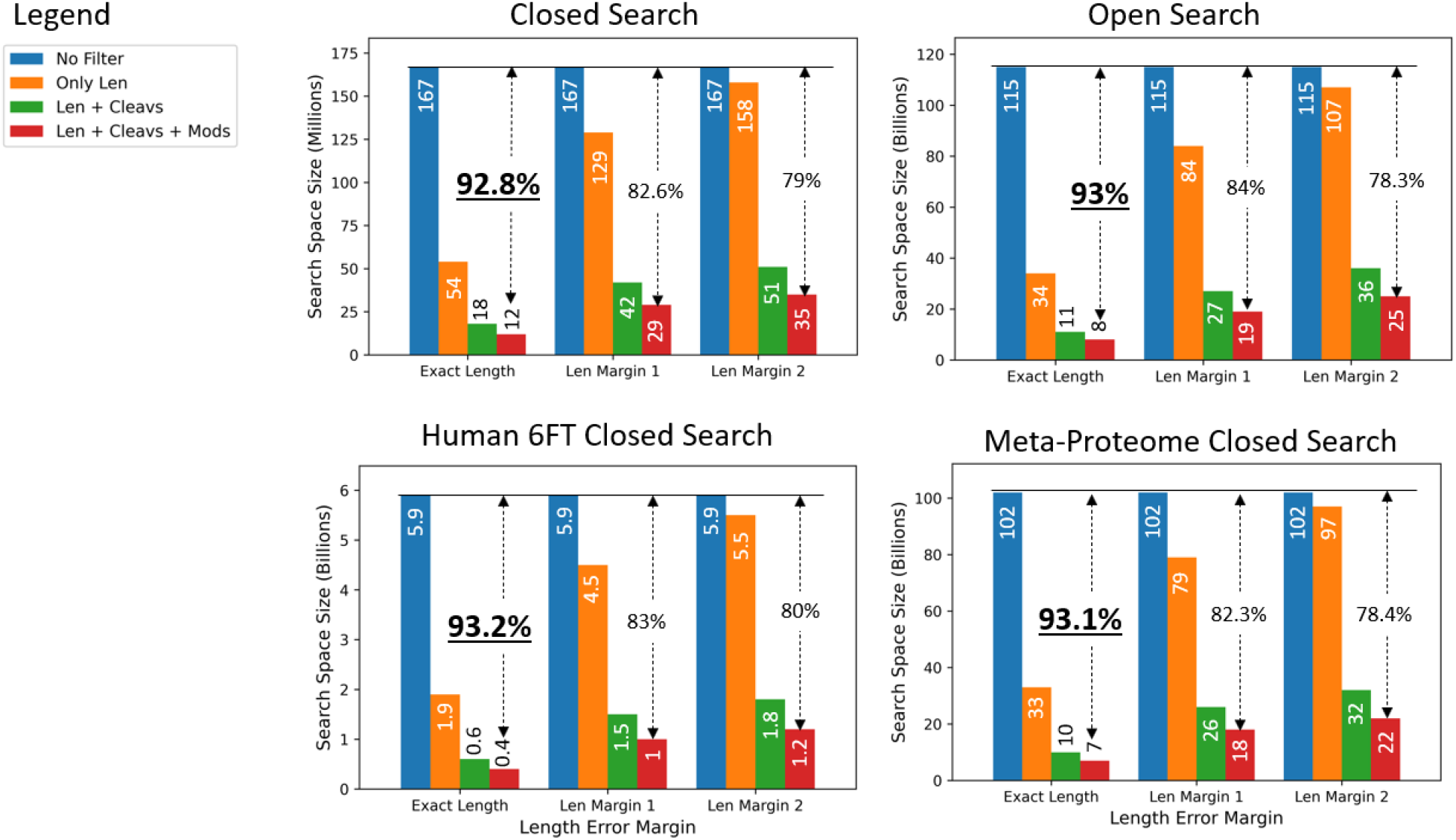
Our preliminary analysis shows that accurately predicting the length, missed cleavages, and modification status can significantly reduce the search space size and provide database search speedup by reducing the number of redundant matches that need to be performed normally. The above analysis is performed for human databases. Three different sets of experiments are performed for different length error tolerances. The speedup gain from the length filter is minimal when allowing error tolerance of more than *±*1.

**Supplementary Figure 2:**
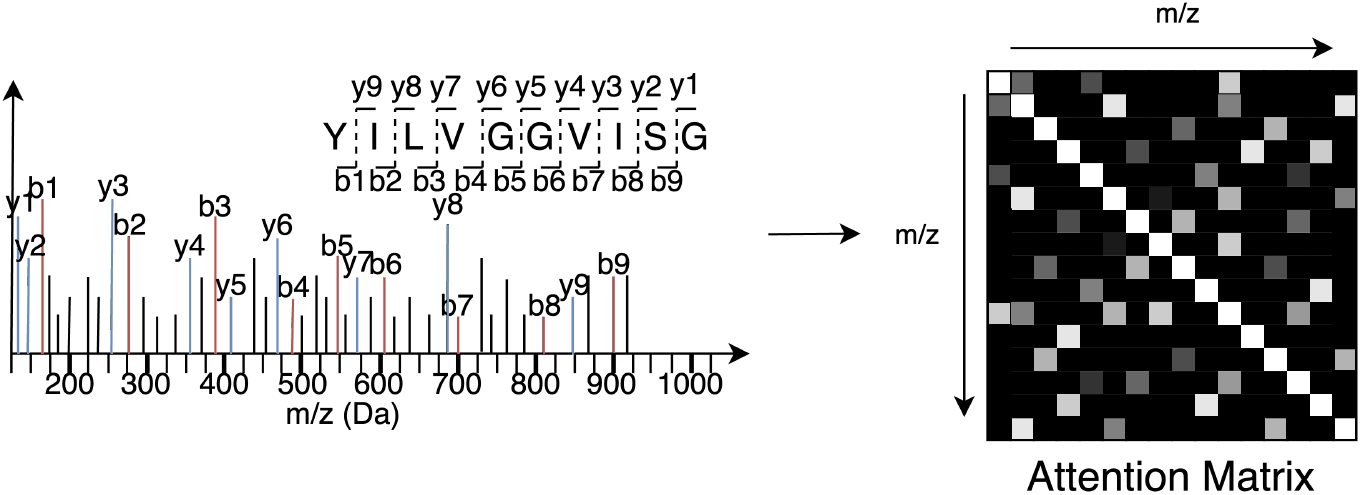
Attention mechanism is useful in capturing long distance as well as short distance relation within a spectrum and allows for more meaningful feature extractions.

**Supplementary Figure 3:**
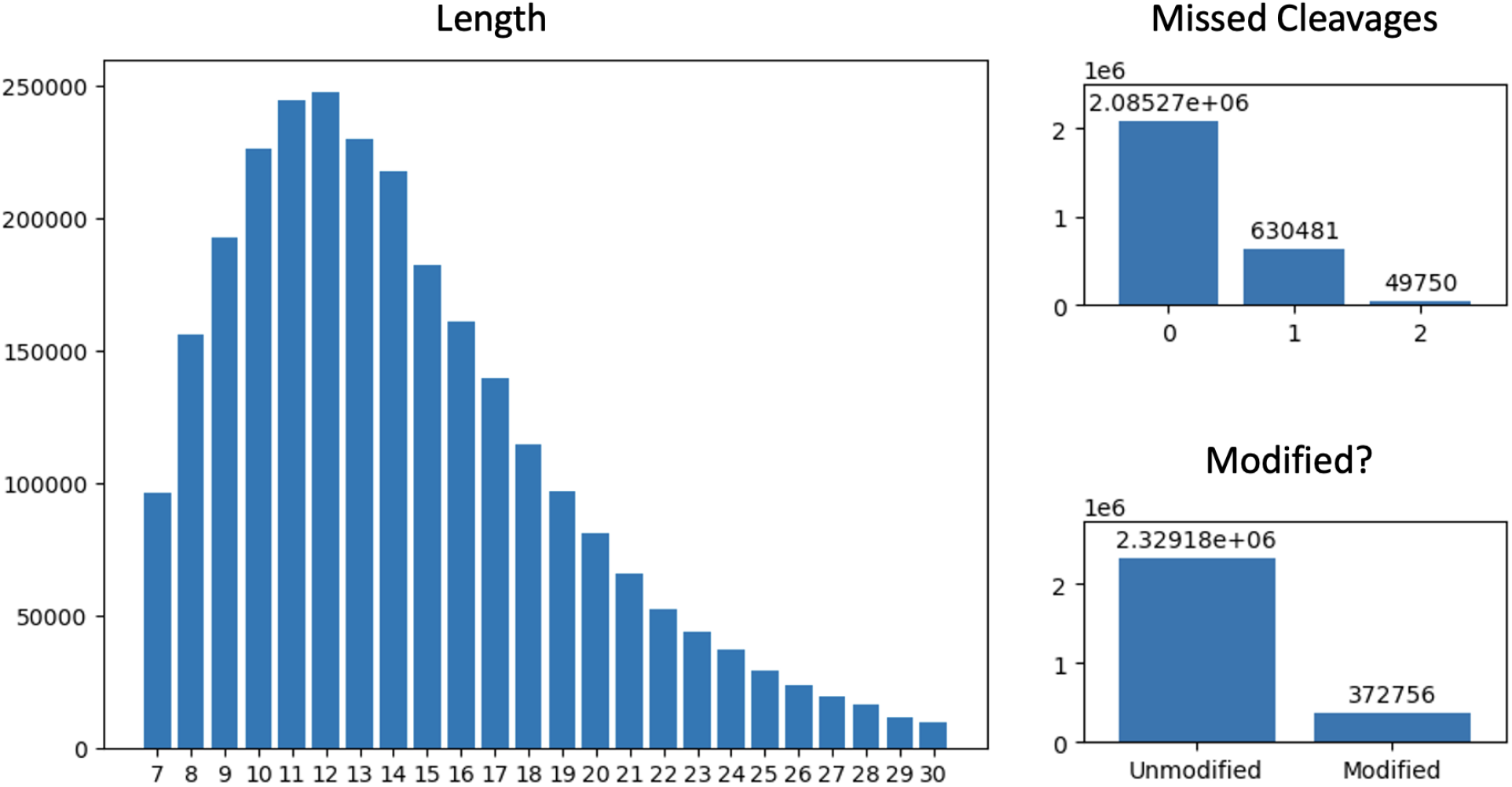
Distribution of peptide length, missed-cleavages, and modifications of training data used in this study. As can be seen the distribution of larger length spectra are fewer in number. To overcome this, we oversample from larger length examples inversely proportional to the count. Similar to length distribution, the examples from 1 and 2 missed cleavages, and examples from modified peptides are oversampled for balancing the training data.

**Supplementary Figure 4:**
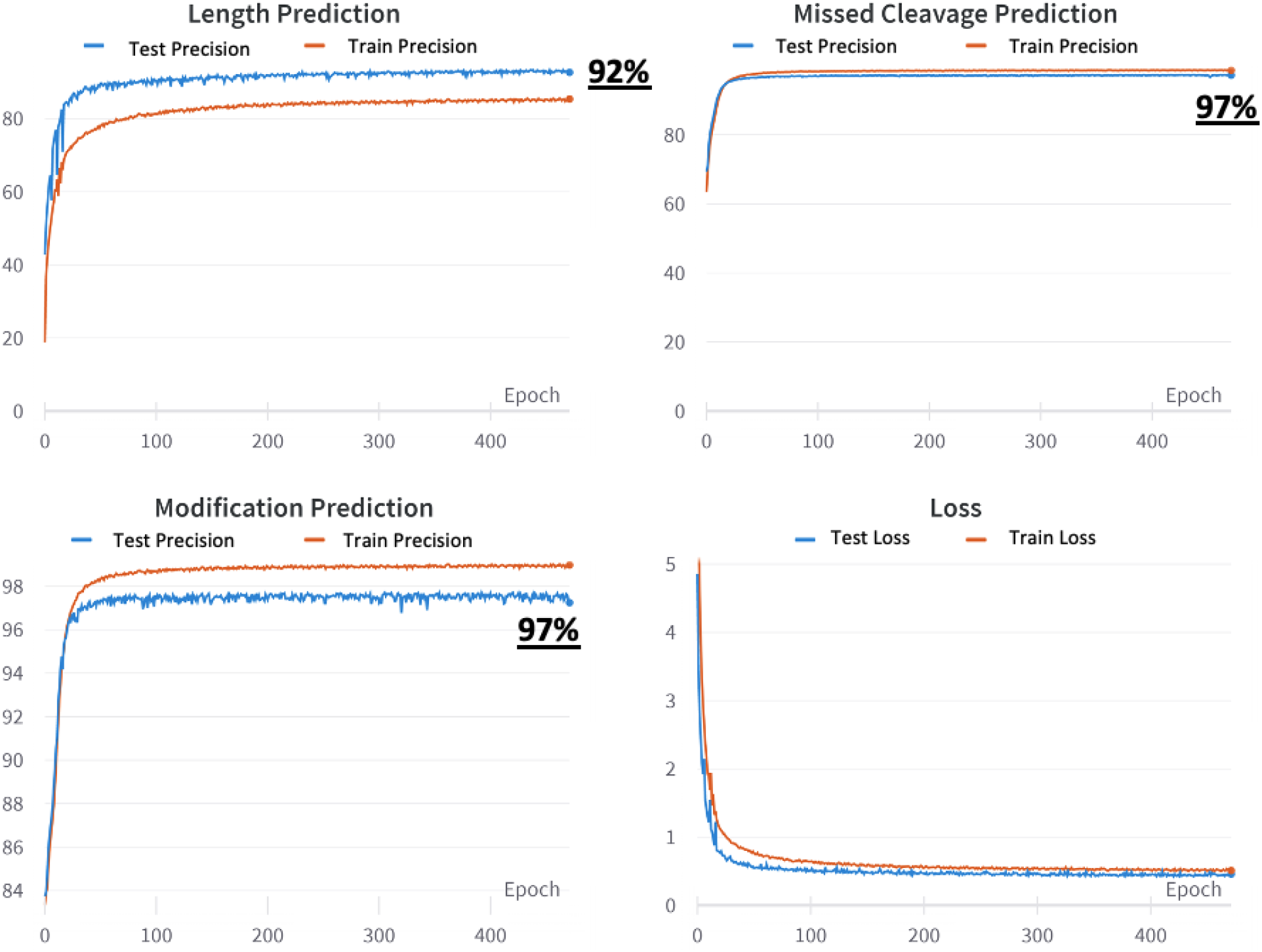
The network is trained for 500 epochs to achieve the test performance of 92% for peptide length, 97% for missed cleavages, and 97% for the status of the modifications.

## Supplementary Tables

**Supplementary Table 1:** Optimal hyperparameters values for GBDT.

| Hyper Parameter | Value |
| --- | --- |
| Max Depth | 30 |
| Min Samples_split | 2 |
| No. of Estimators | 200 |

**Supplementary Table 2:** Optimal GBDT hyperparameters value for Epistemic Uncertainty.

| Hyper Parameter | Value |
| --- | --- |
| Max Depth | 5 |
| Min Samples_split | 2 |
| No. of Estimators | 200 |

